# Observing fear enables plasticity of the dmPFC-BLA pathway during subsequent learning of inhibitory avoidance

**DOI:** 10.1101/296681

**Authors:** Wataru Ito, Alexei Morozov

## Abstract

Observing fear in others is a form of social distress. In mice, it leads to the formation of silent synapses in the prefrontal-amygdala pathway and enhances inhibitory avoidance learning at a later time. Here, we report that observing fear enabled ex vivo LTP. The unsilencing of silent synapses was the prevailing mechanism and the plasticity was occluded by inhibitory avoidance training. In free moving mice with OF experience, the inhibitory avoidance training transiently facilitated local field potentials evoked in BLA by stimulating the prefrontal afferents. The facilitation persisted during the first 4 hours after IA training, which is the time when IA memories consolidate. Thus, the OF-generated silent synapses enable plasticity that may enhance consolidation of IA memories.

## Introduction

Observing how others are expressing fear or pain is a form of psychosocial distress, which can lead to PTSD in humans (*1–5*). The relevant behavioral effects in rodents exposed to distressed conspecifics include formation of contextual and cued fear memories in the observational fear (OF) and fear conditioning by proxy paradigms (*6–8*), enhanced performance in in the two-way avoidance and contextual fear learning tasks performed immediately after a home cage interaction with the demonstrator that just underwent contextual fear conditioning (*9*) and development of anhedonia and anxiety-like behaviors after repeated observation of social defeat in another animal (*10, 11*).

We have recently found that exposure to a conspecific receiving electrical footshocks in the OF mouse paradigm enhances inhibitory avoidance (IA) learning and increases the proportion of the NMDA receptor only (silent) synapses in the input from the dorsomedial prefrontal cortex to the basolateral amygdala one day after OF whereas sub anesthetic ketamine administered immediately after OF abolishes both effects (*12*). It raises a question whether the OF-generated silent synapses participate in the OF-enhanced IA learning. Because silent synapses can incorporate AMPAR and convert to functional synapses (*13–18*), we hypothesized that the OF-generated silent synapses enhance or enable facilitation of the dmPFC-BLA during IA learning. The hypothesis was tested here by quantifying plasticity and silent synapses along the OF-IA paradigm *ex vivo* and by monitoring local field potentials evoked in BLA by stimulating the dmPFC afferents within BLA *in vivo*. OF enabled facilitation of dmPFC-BLA pathway by IA training via unsilencing of the OF-generated silent synapses.

## Materials and Methods

### Animals

129SvEv/C57BL/6N F1 hybrid male mice were produced from breeding trios of one C57BL/6N male and two 129SvEv females, weaned at p21, and housed two littermates per cage as described (*12*). ChR2-AAV virus was injected at p28 and experiments were performed at p60-75, according to Virginia Tech IACUC-approved protocol.

### Behavioral treatments

#### Observational fear

Observational fear (OF) procedure (*6*) was performed in a fear conditioning chamber (Med Associates, St. Albans, VT), divided into two unequal compartments by a transparent Plexiglas wall with 7 mm diameter holes, spaced at 2 cm interval (Fig. 1B), to allow auditory and olfactory cues and whisker to whisker interaction. In the larger 26x20x26 cm compartment, a stainless steel rod floor was covered with a white plastic sheet. In the smaller 26x9x26 cm compartment, the rod floor was exposed. Cagemates observer and demonstrator were placed, respectively, in the larger and the smaller compartments. After 5 min acclimation, 24 footshocks (1 mA, 2 s, every 10 s) were delivered to the demonstrator. The observer was returned to the home cage and housed alone. In the control procedure, demonstrators did not receive footshocks.

**Figure 1.**
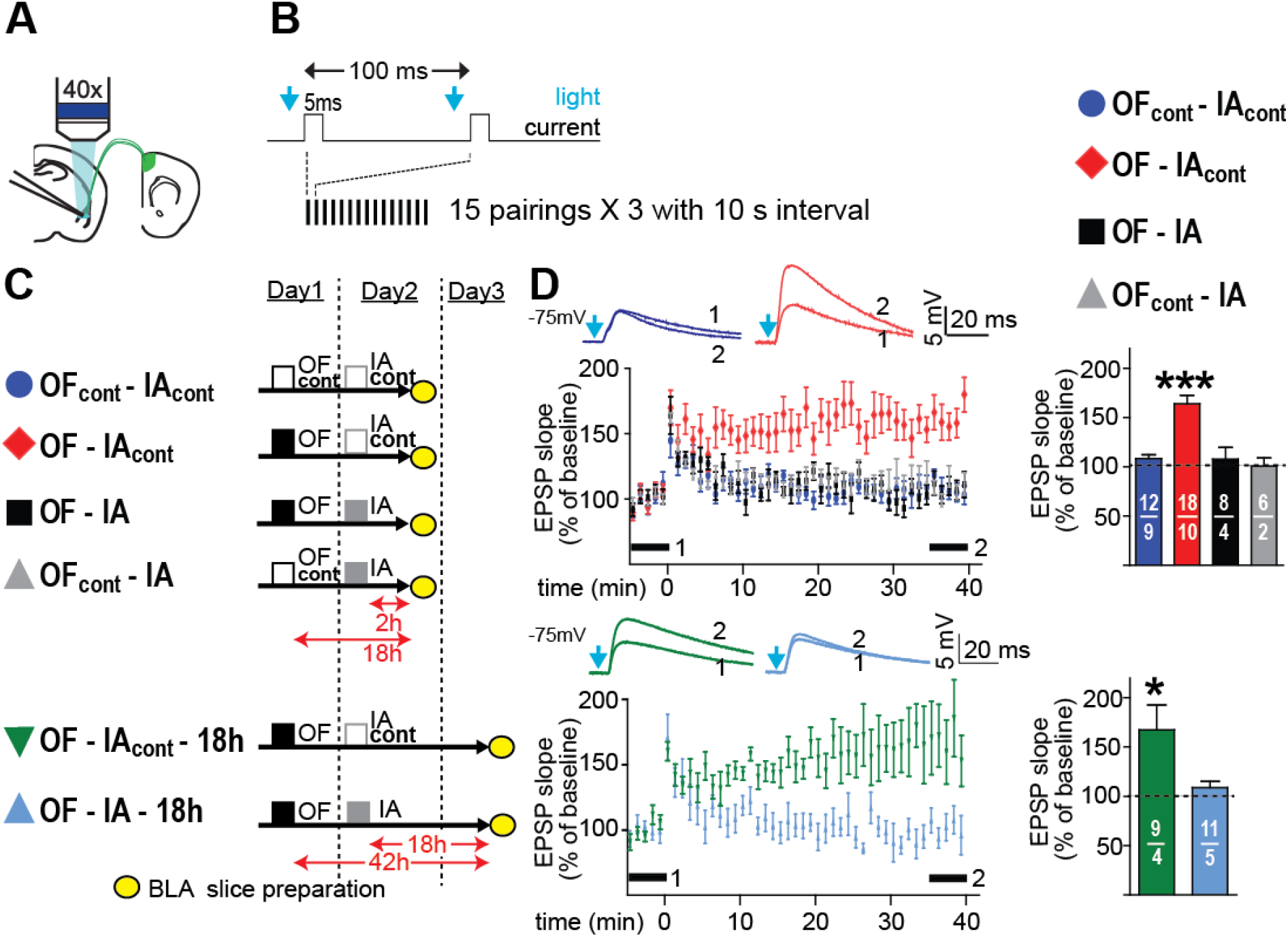
OF enables induction of LTP but IA training occludes it. (A) Recording scheme. (B) LTP induction protocol. (C) Timelines for 6 groups of mice. (D) LTP in dmPFC-BLA pathway. Insets represent examples of averaged EPSPs before (1) and after (2) induction as indicated by thick bars. Right, averaged EPSP slopes at (2). *p<0.05, ***p<0.001, compared to baseline. Number of cells (upper) and animals (lower) are shown on bars here and thereafter.

#### Passive avoidance

The one trial step-through paradigm (*19*) was done in the Gemini Avoidance System with mouse passive avoidance start boxes (San Diego Instruments, San Diego, CA). For training, a mouse was placed in the starting side of the chamber with a guillotine door closed, and lights turned off. After 60 s acclimation, room- and cue-lamps were turned on in the starting side, and the door was opened. Once mice entered the dark side, the door was closed and a 0.15 mA 1 sec electrical footshock was delivered. Mice were returned to home cages 30 s after the door closure. One day later, the same procedure, but without footshock, was used for testing avoidance memory by measuring the latency to enter the dark side.

### In vivo recordings

Mice transduced with AAV-ChR2 in dmPFC and were implanted in BLA with custom made optrodes consisted of a miniature LED coupled to an optical fiber and two 14 uM tungsten wires. LFPs were evoked by 0.5 ms blue light pulses (0.1-2 mW) adjusted to obtain LFP slope at 30-40% of the maximum. LFP were recorded using the RHD2000 series 8 channel evaluation system from Intan Technologies. The data were processed using a custom Matlab script and the Clampfit software. Animals were habituated to the recording setup during two 6 h daily recording sessions or until LFP responses stabilized. The baseline responses were recorded for at least 1 h prior to IA training from the mouse in the home cage without lid. Then, the mouse was disconnected from the recording setup, trained in IA and reconnected to continue the recording for another 6 hours.

### Surgery

ChR2-AAV pseudo-type 1 virus at the titer of 10^12^ viral particles/ml was prepared by University of North Carolina Gene Therapy Vector Core (Chapel Hill, NC) using Addgene plasmid 20071 (*20*), in which ChR2-Venus expression is driven by CAG promoter. P28 mice were anesthetized with Ketamine/Xylazine/Acepromazine (100/20/3 mg/kg) injected intramuscularly after induction by 5% Isoflurane and placed in a stereotaxic apparatus (David Kopf, Tujunga, CA). After minimum craniotomy (~0.5 mm diameters), preserving the dura mater, a heater-pulled glass pipette (shaft: 0.6/0.4 mm external/internal diameter, tip: 50 µm, Drummond, Broomall, PA) filled with the virus solution was slowly lowered to target brain areas. 0.5 µl of the solution were injected bilaterally, using a syringe pump connected to the pipette through plastic tubing filled with water, in dmPFC at 1.3 mm anterior, 0.4 mm lateral from bregma, and 1.3 mm ventral from brain surface or in the temporal association cortex (TeA) at 2.8 mm posterior, 4.6 mm lateral from bregma, and 1.8 mm ventral from brain surface, as described (*21*). For *in vivo* recordings, in addition to the viral transduction of dmPFC, the optrodes were implanted in BLA. For post-operation analgesia, ketoprofen (5 mg/kg) was administered subcutaneously. To prevent body weight loss caused by surgery, Jell-O based calorie supplement was given for one week (4g/animal/day).

### Ex vivo electrophysiology

#### General

Amygdala slices, 300 µm thick, were prepared as described (*21*) using ice cold partial sucrose artificial cerebrospinal fluid (ACSF) solution containing (in mM) 80 NaCl, 3.5 KCl, 4.5 MgSO_4_, 0.5 CaCl_2_, 1.25 H_2_PO_4_, 25 NaHCO_3_, 10 glucose, and 90 sucrose equilibrated with 95 % O_2_ / 5 % CO_2_ (*22*), incubated at 33 °C for 30 min and stored in the same solution at room temperature for at least 45 min before recording. Recording chamber was superfused at 2 ml/min with ACSF equilibrated with 95 % O_2_ / 5 % CO_2_ and containing (in mM) 119 NaCl, 2.5 KCl, 1 MgSO_4_, 2.5 CaCl_2_, 1.25 H_2_PO_4_, 26 NaHCO_3_, 10 glucose and 100 µM picrotoxin (pH 7.4). Whole cell recordings were obtained at 30 ± 1 °C with EPC-10 amplifier and Pulse v8.76 software (HEKA Elektronik, Lambreht/Pfaltz, Germany). Putative glutamatergic neurons in BLA were identified by their pyramidal morphology (*23*) under Dodt gradient contrast optics (cusom made) at 850 nm LED illumination (Thorlabs, Newton, NJ) and were recorded using 2-4 MΩ pipette filled with (in mM) 120 K-gluconate, 5 NaCl, 1 MgCl_2_, 10 HEPES, and 0.2 EGTA, 2 ATP-Mg, 0.1 GTP-Na, for current clamp experiments, or with 120 Cs-methanesulfonate, 5 NaCl, 1 MgCl_2_, 10 HEPES, and 0.2 EGTA, 2 ATP-Mg, 0.1 GTP-Na and 10 mM QX314, for voltage clamp experiments, pH 7.3, osmolarity 285 Osm. Membrane potentials were corrected by the junction potential of 12 mV. Series resistance (Rs) was 10 – 20 MΩ and monitored throughout experiments. Data were not included in the analysis if Rs changed more than 20%. 470 nm 1 ms light pulses were generated using an LED lamp (Thorlabs) and a custom LED driver based on MOSFET and were delivered through a 40× objective lens (Olympus, Center Valley, PA) at 0.3 to 2.5 mW, calibrated by a photodiode power sensor (Thorlabs) at the tip of the lens.

#### AMPAR/NMDAR ratio

AMPAR currents were measured at the peak response in cells held at -75 mV membrane potential. NMDAR currents were determined at 50 ms after the peak in cells held at +40 mV. Stimulus intensity was adjusted to keep the amplitude of AMPAR currents within the range of -50 to -200 pA. AMPAR/NMDAR indices were determined from the averages of 10 to 25 EPSCs.

#### Silent synapses

were quantified by the minimal-like stimulation technique (*24–26*) using the 25 µm diameter light spot. AMPAR EPSC were recorded at -75 mV and mixed AMPAR/NMDAR EPSC - at +40 mV. Stimulus intensity was adjusted to obtain success rate for AMPAR EPSC within the range of 30-70%. No threshold or amplitude criteria of minimal stimulation were applied. Twenty five to 100 traces were recorded at each membrane potential. Failures and responses at -75 mV (F_-75_) and at +40 mV (F_+40_) were identified by visual analysis of traces. The fraction of silent synapses was calculated using an equation: 1- Ln (F_-75_)/Ln(F_+40_) (*27*).

### Data Analysis

Statistical analyses were performed using GraphPad Prism 5 (GraphPad Software, La Jolla, CA). Differences were tested using the two-tale unpaired or paired t-test as indicated and deemed significant with p<0.05.

## Results

### OF enables facilitation of dmPFC-BLA pathway ex vivo whereas IA training reverses this effect

To study synaptic transmission in the dmPFC-BLA pathway, dmPFC was transduced with a ChR2-AAV and synaptic responses to blue light stimulation of the dmPFC axons in BLA were recorded from BLA principal neurons (Fig. 1A). Synaptic plasticity was tested by inducing long-term potentiation (LTP) using a modified spike-timing dependent plasticity (STDP) protocol, in which the onset of excitatory postsynaptic potential (EPSP) preceded action potential (AP) by 8-10 ms (*28*). In our earlier study, this protocol with 75 EPSP-AP pairings has been shown to potentiate the dmPFC-BLA synapses (*21*). Here, to detect possible enhancement of LTP by OF, the number of EPSP-AP pairings was set at 45 (Fig. 1B), which was below the threshold for LTP induction in control mice. Amygdala slices were prepared from mice which had one of the 4 patterns of behavioral experiences (Fig. 1C). The animals underwent OF (“OF”) or the OF control (“OFcont”) procedures followed, one day later, by IA training (“IA”) or the IA training control (IAcont”) procedures and then sacrificed 2 or 18 h later. LTP data are shown in Fig. 1D. LTP was observed in the “OF-IAcont” group (LTP as % of baseline EPSP slope at 35 − 40 min after induction, 164 ± 8.6 %, n = 18, p < 0.0001), but not in the “OFcont - IAcont” group (108 ± 4.1 %, n = 12, p> 0.05), indicating that OF enabled LTP. The “OF-IA” group showed no LTP (109 ± 6.2 %, n = 8, p> 0.05), indicating that IA training abolished the OF-enabled LTP. There was no differences between the “OFcont - IAcont” and “OFcont - IA” groups, indicating that IA training per se did not cause depressing responses to the LTP induction protocol. The “OF-IAcont - 18h” group showed LTP (168 ± 25 %, n = 11, p= 0.03), indicating that the OF-enabled plasticity persisted for 42 h after OF whereas the “OF-IA - 18h” group showed no LTP (108 ± 12 %, n = 6, p> 0.05), indicating that the elimination of plasticity by IA training was persistent as well.

### OF enables postsynaptic mechanisms of plasticity

Several studies provided evidence that LTP in the cortical inputs to the lateral amygdala is expressed presynaptically (*29–35*). To examine LTP expression site in the dmPFC-BLA pathway, we measured the paired-pulse ratio (PPR), an indicator of a change in the presynaptic release probability (*36*), before and after LTP induction in a subset of slices from the OF - IAcont group. The changes in PPR varied among cells and there was a tendency towards a decrease, but it did not reach a statistical significance (baseline, 1.54±0.17; LTP, 1.34±0.17, n=7, p=0.42) (Fig. 2B). The correlation between LTP and CV^-2^, an indicator of presynaptic LTP (*37*), was non-significant as well (Fig. 2C). These data indicated that presynaptic terminals were not the major expression site for the OF-enabled LTP, despite the evidence that LTP in the cortical inputs to the lateral amygdala in naive rodents is expressed presynaptically (*29–35*). One possible reason for the weakness of presynaptic expression in our system is the presence of ChR2, which increases the presynaptic release probability (*38*) and may abolish presynaptic plasticity via the ceiling effect. We studied this by inducing LTP in the slices from naive mice with a stronger STDP protocol (75 EPSP-spike pairings) (*21*), which produced LTP of a comparable magnitude. In contrast to the OF-enabled LTP induced by a weaker protocol (Fig. 2D), the decreases in PPR (baseline: 1.47±0.25; LTP: 1.22±0.23, n=7, p=0.002) (Fig. 2E) and the correlation between LTP and CV^-2^ were significant (Fig. 2F), indicating presynaptic LTP expression. Thus, the presence of ChR2 does not abolish the presynaptic mechanisms of LTP expression.

**Figure 2.**
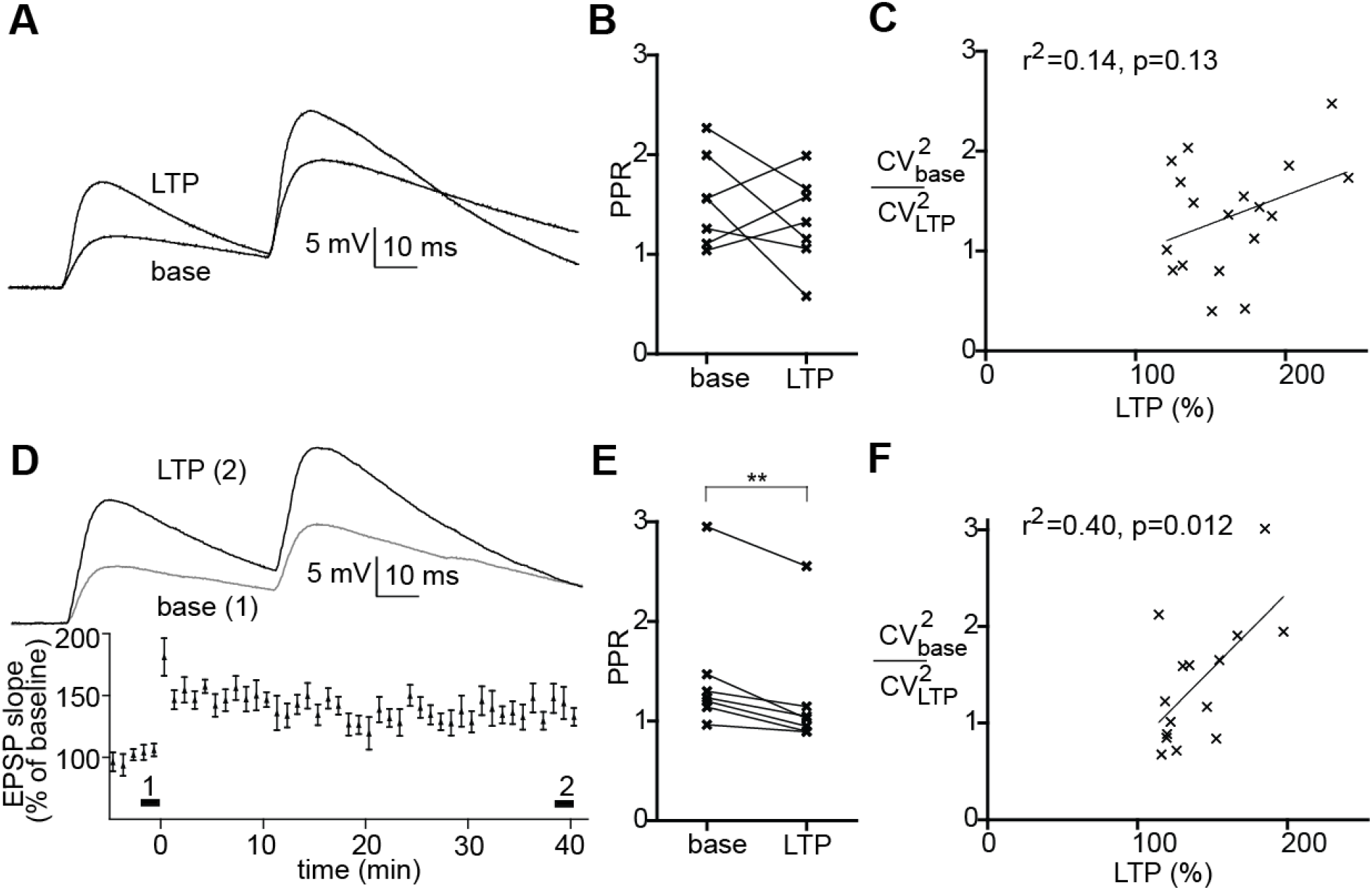
OF-enable plasticity is not expressed presynaptically. (A-C) LTP experiment using the weak STDP protocol in the OF slices. (A) Examples of paired-pulse recording before and after LTP induction. (B) Summary data for paired pulse ratios. (C) Relative CV^-2^ as a function of LTP. (D-F) LTP experiment using the strong STDP protocol in slices from mice that did not undergo OF. (D) Examples of paired-pulse recording before and after LTP induction. (E) Summary data for paired pulse ratios. (F) Relative CV^-2^ as a function of LTP. **p<0.01.

The lack of significant changes in PPR and in the correlation between CV^-2^ and LTP suggest that the plasticity enabled by OF expressed postsynaptically, which was expected to increase the ratio between postsynaptic currents mediated by AMPA and NMDA receptors. Measuring this ratio requires the Cs+-based internal solution, which is not suitable for LTP induction by STDP protocols. As an alternative, we studied a facilitation induced by pairing 80 pulses of the 2 Hz presynaptic stimulation with a steady postsynaptic depolarization to 0 mV (*37*). Because of poor stability of the whole cell patch when membrane potential is alternated between -75 mV for recording the AMPAR EPSCs and +40 mV for the mixed AMPAR+NMDAR EPSCs, the recording duration was limited to about 13 minutes after the induction of plasticity. The pairing protocol induced significant increases in the AMPAR EPSCs in the OF-IAcont group (p=0.0008, n=14), but not in the OFcont-IAcont (p=0.31, n=12) or OF-IA groups (p=0.44, n=15) (Fig. 3B-D). There were no significant changes in the NMDAR EPSCs (Fig. 3E) but an increase in the AMPAR/NMDAR EPSCs ratio in the OF-IAcont group (p=0.02, n=14) (Fig. 3F). These data indicated that OF enabled a postsynaptically expressed plasticity.

**Figure 3.**
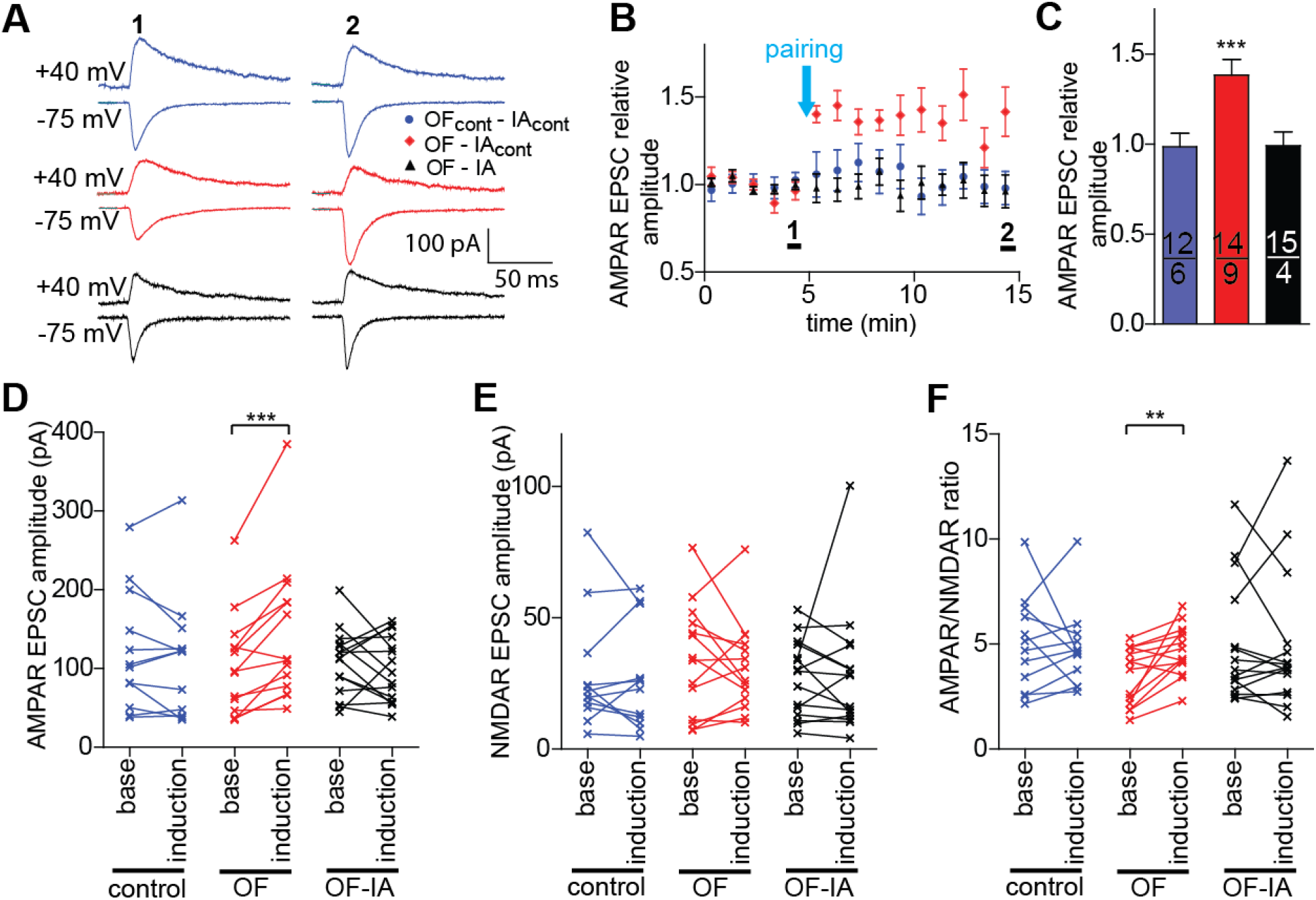
Pairing-induced plasticity in the OF group is mediated by increases in the AMPAR current, but not NMDAR current. (A) Examples of AMPAR (lower) and AMPAR/NMDAR - mixed currents (upper) before (1) and after (2) the pairing in neurons from OF controls (blue), OF (red) and OF-IA (black) groups. Averages of 5-10 consecutive sweeps (grey) are shown. (B) Summary data for the AMPAR EPSC amplitudes before and after the pairing. (C) AMPAR EPSC amplitudes during the last 5 sweeps of recording, normalized to the amplitudes during the last 5 sweeps before the pairing. (D-F) Mean amplitudes of AMPAR EPSCs (D), NMDAR EPSC (E) and their ratios (F) during base line recording (min 0-5) and during the last 5 min (min 10-15).

### Presence of silent synapses coincides with the OF-enabled plasticity and is abolished by IA training

We showed earlier that OF generated silent, NMDAR-only, synapses, detectable in the dmPFC pathway at about 18 h after OF (*12*). Here, we used the minimum-like stimulation method to quantify silent synapses in slices prepared at different time intervals after OF, ranging from 4 h to 1 week (Fig. 4A). There was a significant presence of silent synapses at 18h and 42h, the same time points after OF when LTP was readily inducible (Fig. 4D, E, I). There were no significant amounts of silent synapses at 4 h and 1 week after OF (Fig. 4 B-F, I). In mice that underwent IA training at 18 h after OF, there were no significant amount of silent synapses at 2 and 18 h after IA training, which indicated that IA training eliminated silent synapses generated by OF and it they did not reappear 18 h later (Fig. 4 G, H, I).

**Figure 4.**
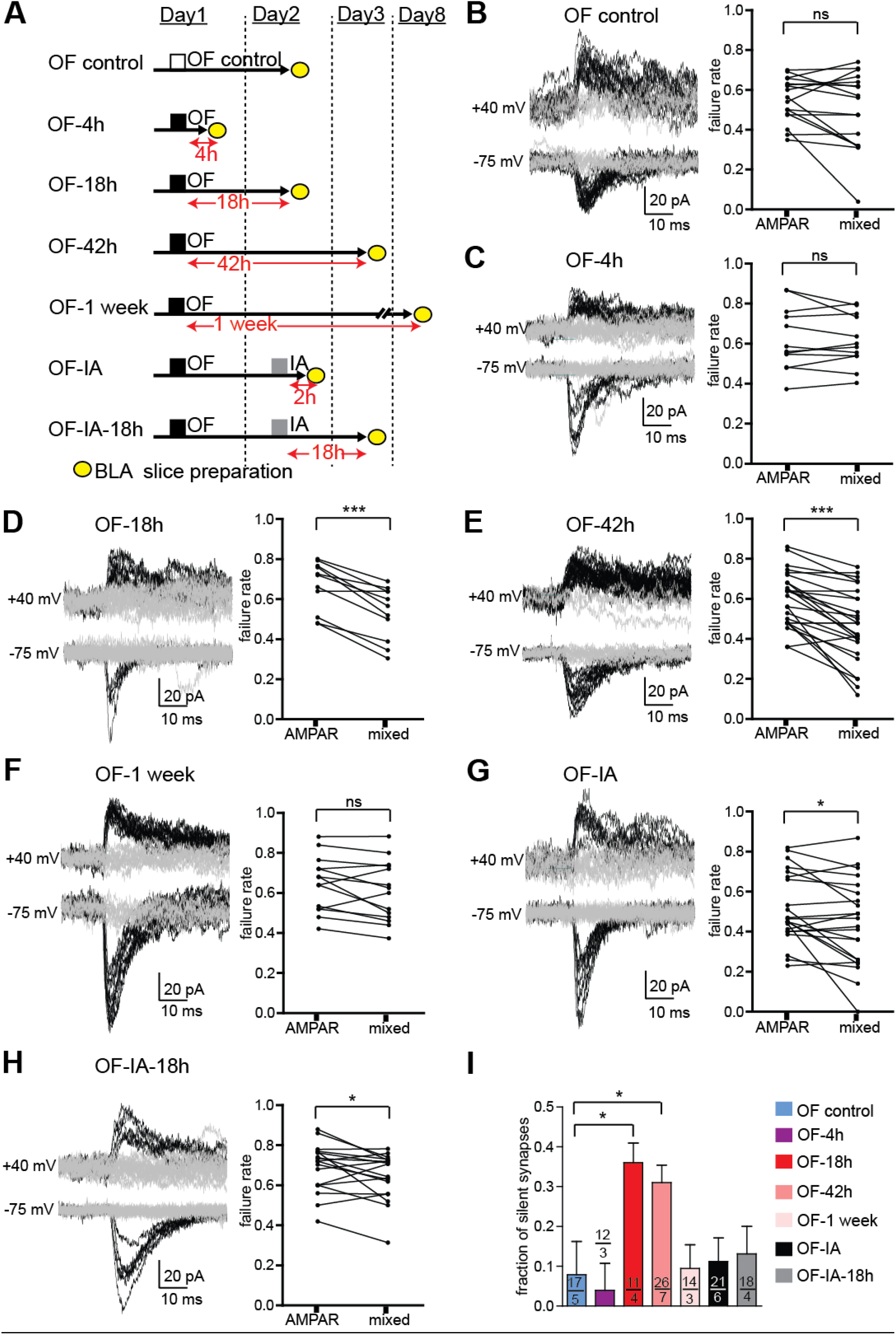
OF generates silent synapses and IA training eliminates them. (A) Seven experimental groups. (B-H) Left: Example traces at -75 (lower) and +40 mV (upper). Successes EPSC responses and failures are shown respectively in black and grey. Right: Failure rates of AMPAR and AMPAR-NMDAR mixed currents and estimated fraction of silent synapses. (H) Summary data for fraction of silent synapses. * p < 0.05.

## Observing fear enables facilitation of dmPFC-BLA pathway by IA training in the free moving mice

To test whether the *ex vivo* plasticity of dmPFC-BLA synapses, elicited by OF and then abolished by IA training, reflects an actual *in vivo* facilitation during IA training, we recorded local field potentials evoked in BLA by blue light stimulation of dmPFC axons expressing ChR2. The mice were transduced with AAV-ChR2 in dmPFC and implanted with optrodes in BLA (Fig. 5A, B). The mice underwent OF or the OF control procedure on and then were trained in IA on the next day. The evoked LFPs were recorded before and after IA training, but not during IA training when animals were disconnected. Once reconnected after IA training, all mice showed decreased LFPs, which we normally observe after handling. At about 40 min after IA training, LFPs in the control mice returned to the baseline, whereas in the OF mice, the LFPs showed a facilitation of about 25%, which lasted for about 4 h (Fig. 5C, D). It indicates that OF enabled an *in vivo* facilitation of the synaptic transmission between dmPFC and BLA by IA training.

**Figure 5.**
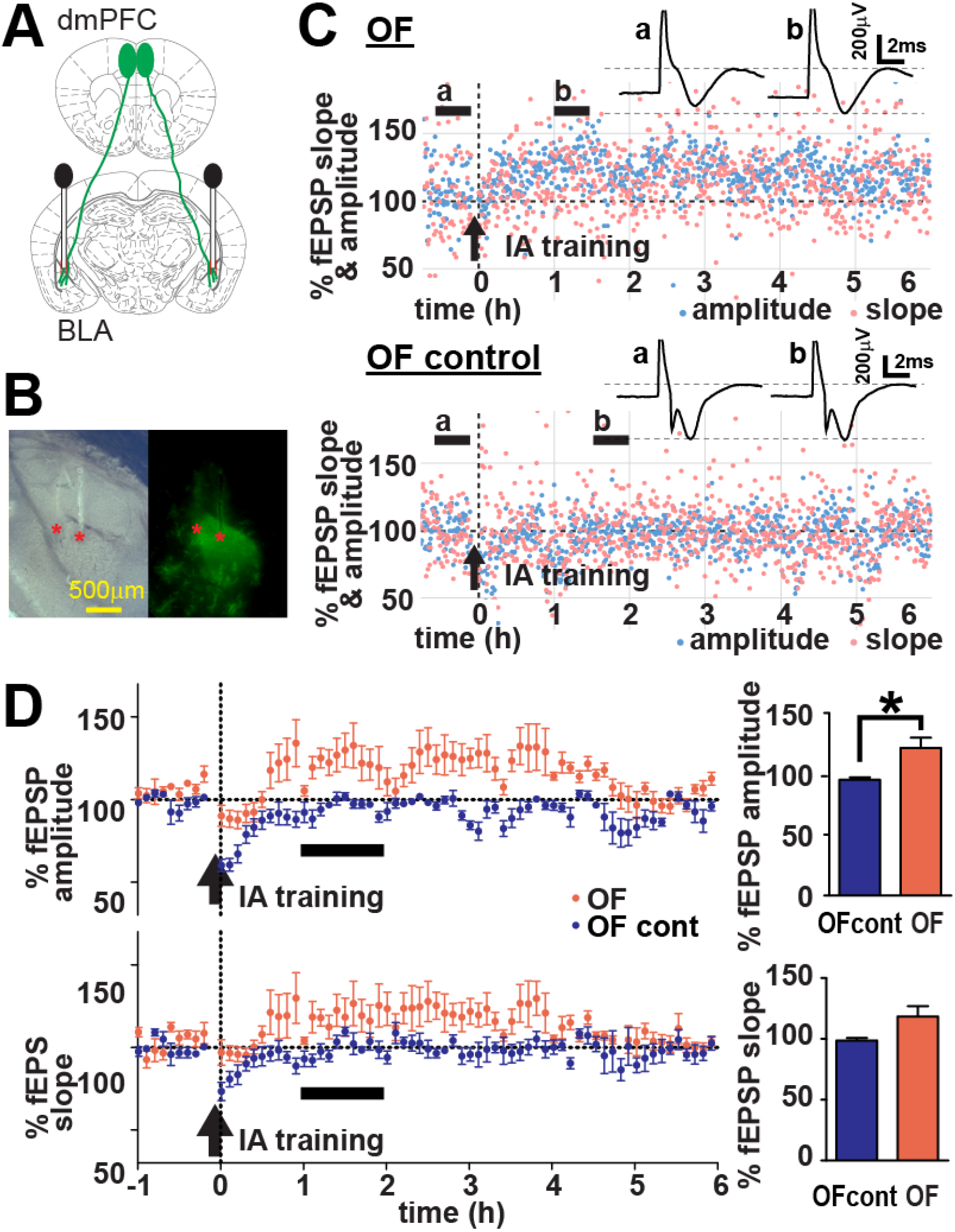
OF enables facilitation of the dmPFC-BLA pathway by IA training in free moving mice. (A) Schematics representing ChR2-Venus expressing dmPFC neurons projecting to BLA and bilaterally implanted optrodes. (B) Verification of optrode location. Left: visible light, right: Venus fluorescence images. Position of electrode tips marked as red stars. (C) (Upper) Examples of fEPSP traces before (a) and after (b) IA training (averaged over the time range shown by horizontal black bars) in an OF-experienced mouse (OF) and an OF control mouse (OF control). (Lower) fEPSP slopes and amplitudes normalized to the baseline for the same animals. (D) Summary fEPSP data. n=8 (OF), 7 (OF cont). Horizontal black bars indicate the time range corresponding to the averaged fEPSP data on the right. Vertical arrows indicate the time of IA training. Each data point represents the average during 6 min. Blue light pulses (3 mW, 0.5 ms duration) were given every 30 s. *p<0.05.

## Discussion

The main finding of this study is that observing fear in others enables plasticity in the dmPFC-BLA synapses both *ex vivo* and *in vivo*.

The ex vivo STDP LTP was tested because it relies on co-incidental presynaptic activation and postsynaptic spiking and may represent a process relevant to associative learning (*28*). The weak STDP protocol, which does not induce LTP in the dmPFC-BLA pathway of naive mice, produced LTP in mice that underwent OF 18 or 42 h before slice preparation. It indicated that neuronal changes that enable such LTP last for at least 2 days after OF. Subjecting mice to IA training 2h or 18h before LTP induction prevented LTP induction in slice, suggesting that the mechanisms of plasticity emerging after OF are depleted by IA training. Such occlusion of LTP by associative learning is a common phenomenon in several brains areas including amygdala and hippocampus (*29, 31, 39*). It indicates that associative learning facilitates the same synapses via a shared mechanism. Our analysis of the paired-pulse ratios, which reflect the presynaptic release probability (*36*), did not show significant differences before and after LTP induction, suggesting that LTP was not expressed presynaptically. It was consistent with the lack of a correlation between LTP and the value of 1/CV2, another indicator of presynaptic LTP expression. The lack of presynaptic LTP was not an artifact of ChR2 expression, which increases the presynaptic release probability (*38*) and can have ceiling effect on presynaptic LTP, because, with a stronger STDP protocol applied to a control group, we obtained LTP with presynaptic expression, judged by the decreases in PPR and strong correlation between LTP and 1/CV2. Since the conditions of STDP experiments were not suitable for isolating the AMPAR- and NMDAR-mediated currents to test the postsynaptic plasticity mechanisms, we employed the pairing plasticity protocol, which allowed measuring AMPAR currents and NMDAR currents before and after the induction of plasticity. The enhancement of AMPAR-currents and not NMDAR currents in the OF group, but not in the OF controls, indicated that OF enabled a post-synaptically expressed plasticity mediated by membrane insertion of AMPAR. Lack of this plasticity after IA training suggests that IA training causes the same AMPAR insertion and abolishes this plasticity in slice. One limitation of this analysis of the pairing-induced facilitation, it did not test for long-lasting changes because of the difficulty of obtaining stable whole cell recording when the holding membrane potential is repeatedly switched between -75 mV and +40 mV. However, it revealed a greater potential for long-lasting postsynaptic changes after OF. The AMPAR insertion is a plausible mechanism to explain the role of silent synapses generated after OF in the observed plasticity (*13*). The coincidental presence of silent synapses and STDP-induced LTP at 18 h and 42 h after OF, and coincidental absence of silent synapses and plasticity in the IA-trained mice suggest that silent synapses are responsible for this LTP.

Our in vivo recording revealed a transient facilitation of LFPs evoked in BLA by stimulation of dmPFC axons. The unsilencing of silent synapses demonstrated ex vivo provides a plausible mechanism for LFP enhancement, but other factors including excitability of postsynaptic neurons and the balance between evoked excitation and inhibition may also contribute to these changes in LFP.

The transient facilitation of amygdala responses to dmPFC input was observed during 4 h after IA, which is the critical period when IA memories consolidate and when consolidation can be enhanced by co-activating BLA, prefrontal cortex and several other BLA-connected areas (*40–42*). The timing of the facilitated BLA responses is a strong indication of their possible role in the OF-enhanced inhibitory avoidance.

## Acknowledgements/Funding Sources

The study was supported by the NIH grant MH112093.

